# Hippocampal *Fos*-expressing neurons stably encode boundaries in novel environments

**DOI:** 10.1101/2025.10.06.680713

**Authors:** Jun Young Oh, Daniel A. Dombeck

## Abstract

The hippocampus recruits place cell ensembles to represent novel environments, but how gene expression relates to this reorganization and its stabilization remains unclear. We imaged calcium transients and *Fos*-expression changes across ~30,000 CA1 neurons and found that neurons with higher *Fos* increases were more likely to be place cells in novel environments, rapidly cluster around environmental boundaries, and maintain stable boundary encoding across days, suggesting a role of *Fos* expression in place cell organization and stabilization.

## Main

Place cell populations in hippocampal CA1 rapidly reorganize in novel environments^1^ and continue to evolve across days through processes known as global remapping^2–4^and representational drift^5,6^, respectively. The *Fos* immediate early gene is a widely used biomarker to label neurons involved in encoding and recall^7–10^ of contextual fear memories, and more recent research is beginning to find links between *Fos*-expression and place coding^11,12^. For example, in familiar environments, higher *Fos-*expression change (increase) was found in stable place cell populations, and lower *Fos*-expression change in reward encoding populations^12^. In novel environments where spatial memories first form, however, it remains unclear how changes in *Fos*-expression relate to place-field location and stability.

To address these questions, we simultaneously imaged calcium transients using the red calcium indicator jRGECO1a^13^ and *Fos-*expression change (*Fos-*change) using the green indicator cfos-shEGFP^8^ in hippocampal CA1 pyramidal neurons (30,900 neurons; 15 mice; 211 sessions) using two-photon microscopy (Fig.1a and Methods). The head-fixed mice were exposed to either novel (N) or familiar (F) virtual environments (VR) with a hidden water reward in the middle of a 3-m track (Fig. 1b, Extended Data Fig. 1a, and Methods). The mice started each discrete trial at the track start (start boundary), received the reward in the middle, and then finished at the track end (end boundary), where the environment froze for 3 seconds before the mice were teleported back to the track start for the next trial. In familiar environment sessions, mice had similar velocity and licking profiles across both early and late laps, with anticipatory slowing and licking before the reward. In novel environment sessions, in contrast, mice reduced their velocity significantly across the track and did not display anticipatory slowing or licking before the reward on early laps (Extended Data Fig 1d-g).

**Fig. 1.**
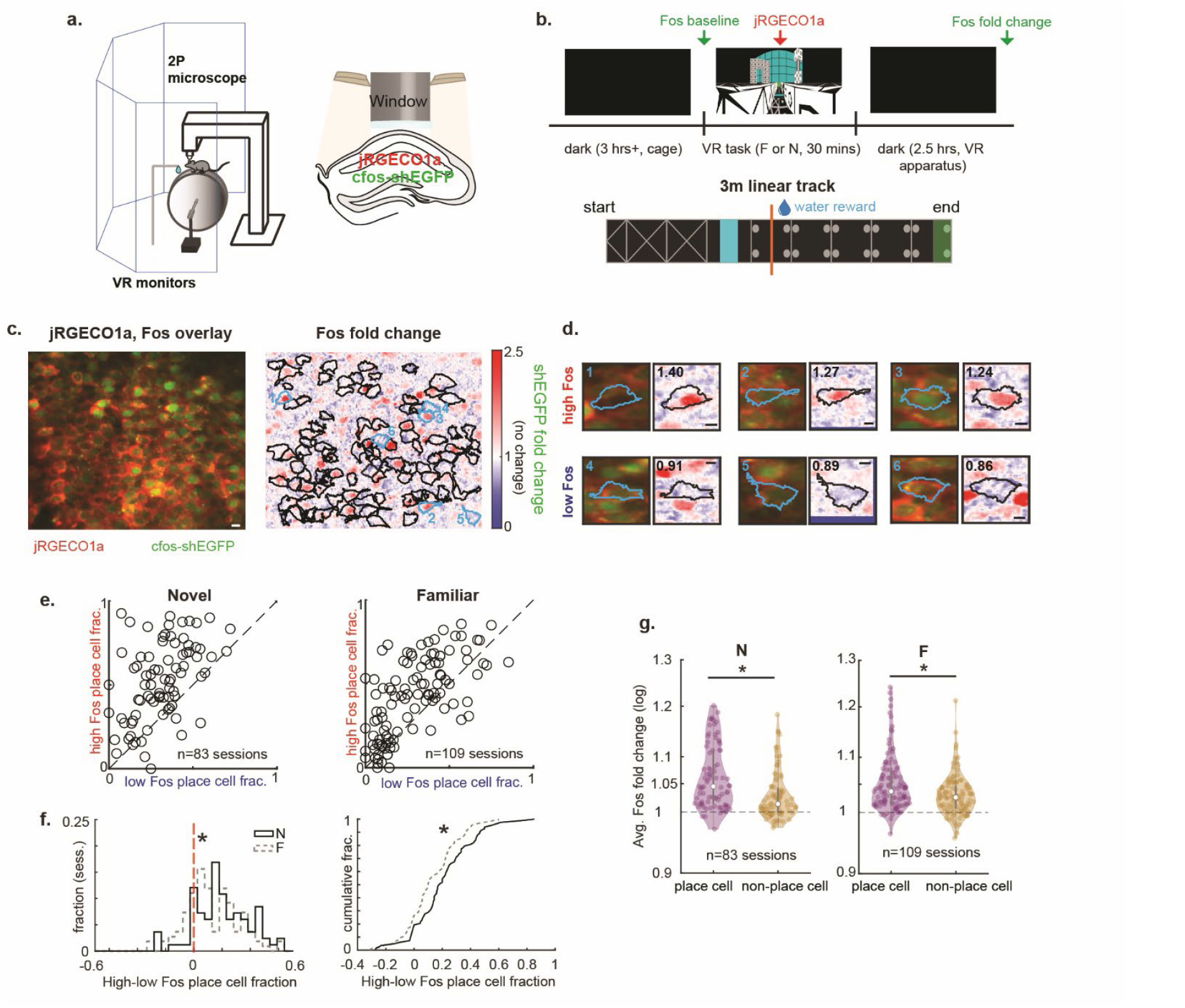
Environmental novelty increases *Fos* change and the *Fos* recruitment of CA1 place cells. **a**. During 2P imaging, head-fixed mice run on a cylindrical treadmill to traverse virtual reality (VR) linear tracks and receive water rewards through a lick spout (left). Hippocampus CA1 is imaged through a cannula window (right). **b**. Experimental design. The mice were placed in enclosed, dark cages with a white noise for at least ~3 hrs, exposed to novel (N) or familiar (F) VR environments for 30 minutes (hidden water reward in the middle of a 3-m track), and then remained in the dark on the treadmill for an additional 2.5 hours. The *Fos* baseline (pre-VR) was imaged with cfos-shEGFP (green), calcium transients during VR with jRGECO1a (red), and *Fos* change was assessed 3h post-initial VR exposure. **c**. Hippocampus CA1 pyramidal cells co-expressing jRGECO1a and cfos-shEGFP. Overlay is shown on the left, and the *Fos* fold change from the baseline is shown on the right with cell ROIs drawn based on calcium transients in VR. Example cells (Fig. **1d**) are highlighted in light blue. The scale bar indicates 10 µm. **d**. Example cells (n=6) with their associated *Fos*-change calculated by averaging shEGFP intensity change relative to baseline within an ROI (black outline). High *Fos*-change cells (top 15% *Fos* change) are shown on top, low *Fos*-change cells (bottom 15% Fos change) on the bottom. The scale bar indicates 5 µm. **e**. For each session, the average fractions of place cells amongst high *Fos*-change cells are plotted against those of low *Fos*-change cells (n=83 sessions from 13 mice for N, 109 sessions from 15 mice for F; two-sided Wilcoxon signed-rank test, high vs low *Fos*-change fraction; F: *P*=4.4e-10; N: *P*=4.9e-11). **f**. Histogram (left) and cumulative distribution function (right) of session-by-session high *Fos*-change cell fraction minus low *Fos*-change cell fraction (n=83 sessions from 13 mice for N, 109 sessions from 15 mice for F; two-sided Wilcoxon rank-sum test, F vs N histogram distribution comparison: *P*=0.007; two-sided, two-sample Kolmogorov–Smirnov test, F vs N cumulative distribution function comparison: *P*=0.007. Star indicates *P*<0.05). **g**. The session-by-session N (left) and F (right) average *Fos* fold change for place cells and non-place cells. Middle values are medians, and the vertical bars are 25^th^ and 75^th^ percentiles (n=83 sessions from 13 mice for N, 109 sessions from 15 mice for F; two-sided Wilcoxon signed-rank test place cell vs non-place cell *Fos* fold change: F: *P*=3.08e-9; N: *P*=1.3e-11; two-sided Wilcoxon rank-sum test; N vs. F place cell vs. non-place cell average *Fos* change difference, *P*=0.018. Star indicates *P*<0.05)

We first categorized cells into high and low *Fos-*change groups based on *Fos* expression change percentile (top and bottom 15^th^ percentile of *Fos-*change within each session, Fig. 1c-d, Extended Data. Fig. 1b and Methods). Across all cells and sessions, *Fos-*change was highest in the novel environment, followed by familiar and dark environments (Extended Data Fig.1c). We then investigated the relationship between *Fos-*change and a cell’s likelihood to be a place cell (Extended Data Fig. 1h) in familiar and novel environments. In familiar environments, consistent with previous research^12^, high *Fos-*change cells were more likely to be place cells and had lower position decoding error compared to low *Fos-*change cells (Fig. 1e,f, Extended Data Fig. 1l). Also, the place cell population as a whole had a higher *Fos-*change compared to non-place cells (n=109 familiar sessions, 15 mice, total 5773 place cells and 10,343 non-place cells; Fig.1g, Extended Data Fig. 1k, Methods). In novel environments, similarly, high *Fos*-change cells were more likely to be place cells with lower position decoding error, and the place cell population had a higher *Fos-*change compared to non-place cells (n=83 novel sessions, 13 mice; total 4794 place cells and 6843 non-place cells; Fig. 1e,g, Extended Data Fig. 1i-j). Notably, these effects were more pronounced in novel environments compared to familiar environments, the difference in place cell and non-place cell *Fos*-change being higher in a novel environment (Fig. 1f-g, Extended Data Fig. 1i). Thus, high *Fos-*change cells are more likely to be place cells than low *Fos-*change cells, and novel environment exposure increases this place encoding difference.

Next, we examined whether the amount of *Fos-*change in place cells was associated with their place code organization (i.e., the location of their place fields). First, to visualize the spatial tuning of high and low *Fos-* change place cells, we sorted and cross-validated heat maps for both novel and familiar sessions (Fig. 2a,d; Extended Data Fig. 2a,c). In both novel and familiar environments, we found that high *Fos-*change place cells clustered preferentially at the track start and end (i.e., track spatial boundaries; Fig. 2a,d,g,h), with *Fos* change being higher at the track boundaries compared to the reward location across all place cells (Fig. 2b,e). Low *Fos* place cells, on the other hand, were distributed between the (post-) reward location and track boundaries in novel environments with a shift to cluster around the (pre- and post-) reward location in familiar environments after learning the environment over days (Fig. 2a,d,g,h, and Methods). Accordingly, high *Fos-* change place cell fractions at the track boundary were higher than those of low *Fos-*change cells, with high *Fos*-change cells’ decoding error being lower at the track boundaries; on the other hand, low *Fos-*change place cells had a greater fraction at the reward location (Fig. 2c,f,i,j, Extended Data Fig. 2b,d). We then analyzed how quickly these different location-encoding preferences formed in the novel environment by dividing the novel environment sessions (30 minutes) into 10 time bins (3 min/each). We found that the high *Fos-*change place cell preference for encoding track boundaries emerged rapidly (first time bin) and remained consistent throughout the session, with a higher boundary fraction than that of low *Fos*-chance place cells (Fig. 2k, Methods). In contrast, the low *Fos-*change (vs high *Fos-*change) place cell preference for encoding the reward zone emerged gradually throughout the session and stabilized in a familiar environment (Fig. 2l). Together, these findings establish that higher *Fos*-change place cells are more likely to encode spatial boundaries in both novel and familiar environments, and this preference for encoding location forms rapidly in novel environments.

**Fig. 2.**
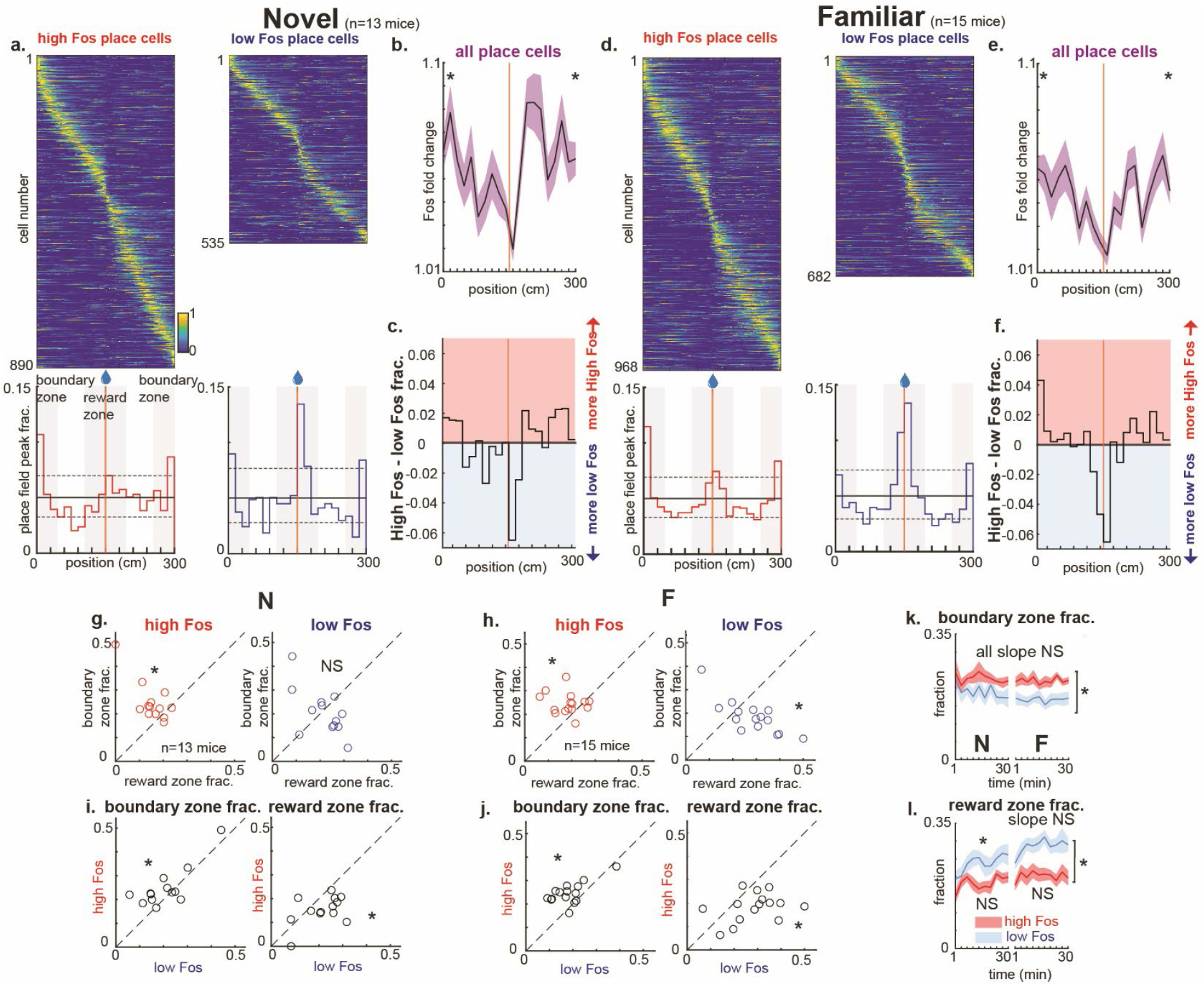
High *Fos*-change place cells rapidly encode spatial boundaries in novel environments. **a-f**. Cross-validated and sorted heatmaps (odd laps sorted by even laps, top) for high *Fos*-change place cells (red) and low *Fos*-change place cells (blue), and their histograms of place field peak fractions (bottom) in N (**a**, n=890 high *Fos* place cells, 535 low *Fos* place cells pooled from 13 mice) and F (**d**, n=968 high *Fos* place cells, 682 low *Fos* place cells pooled from 15 mice). The solid horizontal lines indicate chance level, and dotted horizontal lines indicate two-sided binomial 99% confidence intervals. The difference (high *Fos* minus low *Fos*) in these histograms’ peak fractions is shown for N (**c**) and F (**f**). The average *Fos* change (solid line) and s.e.m. (shaded region) across the track positions for all place cells is shown for N (**b**) and F (**e**) (n=4794 place cells distributed across track positions for N, n=5774 place cells in F; two-sided Wilcoxon signed-rank test; total 20 position bins, N boundary bins (1-4, 17-20) vs. reward bins (9-12): *P*=4.6e-7, F boundary bins vs. reward bins: *P*=4.3e-13. Star indicates *P*<0.05. **g-h**. Fractions of high or low *Fos*-change place cells in the boundary zone plotted against those in the reward zone for N (**g**) and F (**h**) (n=13 mice for N, 15 mice for F; two-sided Wilcoxon signed-rank test, boundary vs reward zone fractions; N-high *Fos*: *P*=0.0061; N-low *Fos*: *P*=0.62, F-high *Fos*: *P*=0.0084; F-low *Fos*: *P*=0.030. Star indicates *P*<0.05, NS indicates statistically not significant). **i-j**. Boundary zone or reward zone fractions of high vs. low *Fos* change place cells for N (**i**) and F (**j**) (n=13 mice for N, 15 mice for F; two-sided Wilcoxon signed-rank test, high vs low *Fos* fractions; N-boundary zone: *P*=0.033; N-reward zone: *P*=0.021, F-boundary zone: *P*=0.0026; F-reward zone: *P*=0.0034. Star indicates *P*<0.05). **k-l**. Boundary zone (**k**) and reward zone (**l**) fractions of high *Fos*-change place cells and low *Fos*-change place cells over time (N on the left, F on the right). Line indicates average, shaded region indicates s.e.m. (n=13 mice for N, 15 mice for F; two-sided Wilcoxon signed-rank test; N-high vs. low *Fos* average boundary zone fraction across 10 time bins: *P*=0.0034, N-high vs. low *Fos* average reward zone fraction: *P*=0.013, F-high vs. low *Fos* average boundary zone fraction: *P*=6.1e-5, F-high vs. low *Fos* average reward zone fraction: *P*=4.3e-4; one sample, two-sided t-test; N-high *Fos* boundary zone slope: *P*=0.57, N-low *Fos* boundary zone slope: *P*=0.076, F-high *Fos* boundary zone slope: *P*=0.95, F-low *Fos* boundary zone slope: *P*=0.92, N-high *Fos* reward zone slope: *P*=0.16, N-low *Fos* reward zone slope: *P*=0.016, F-high *Fos* reward zone slope: *P*=0.61, F-low *Fos* reward zone slope: *P*=0.10. Star indicates *P*<0.05, NS indicates statistically not significant).

Lastly, motivated by findings in familiar environments that *Fos* stabilizes the spatial code^12^, we explored whether the novelty-evoked increase in *Fos* expression in a novel environment predicted the stability of spatial coding of CA1 neurons across days. We tracked the same CA1 cells over five days, imaging them on the first day in a novel environment (day 1), training the mice for three subsequent days in the same environment (days 2-4) and then imaging those cells again in the (now familiar) environment on day 5 (n=30 day 1-5 sessions across 10 mice; Fig. 3a). We utilized our recently developed volumetric registration method to identify the same imaging plane across days with high precision^6^ (Extended Data Fig. 3a-b, and Methods). We categorized CA1 cells as high or low *Fos*-change cells based on day 1 (same method as Fig. 1), and found that day 5 *Fos* change was higher in the day 1 high *Fos* group than in the day 1 low *Fos* group (Extended Data Fig.3c). In addition, multiple representational drift metrics revealed that high *Fos*-change cells overall exhibited across-day representational stability comparable to that of low *Fos*-change cells as the novel environment became familiar (Fig. 3b-d, Extended Data Fig. 3d). However, when we inspected the stability of high and low *Fos*-change populations as a function of track position, high *Fos*-change cells consistently demonstrated significantly higher stability near the boundaries (Fig.3b-i, and Methods). Therefore, higher *Fos*-change in neurons during spatial memory formation predicts an increased likelihood of stable spatial encoding around spatial boundaries across days.

**Fig. 3.**
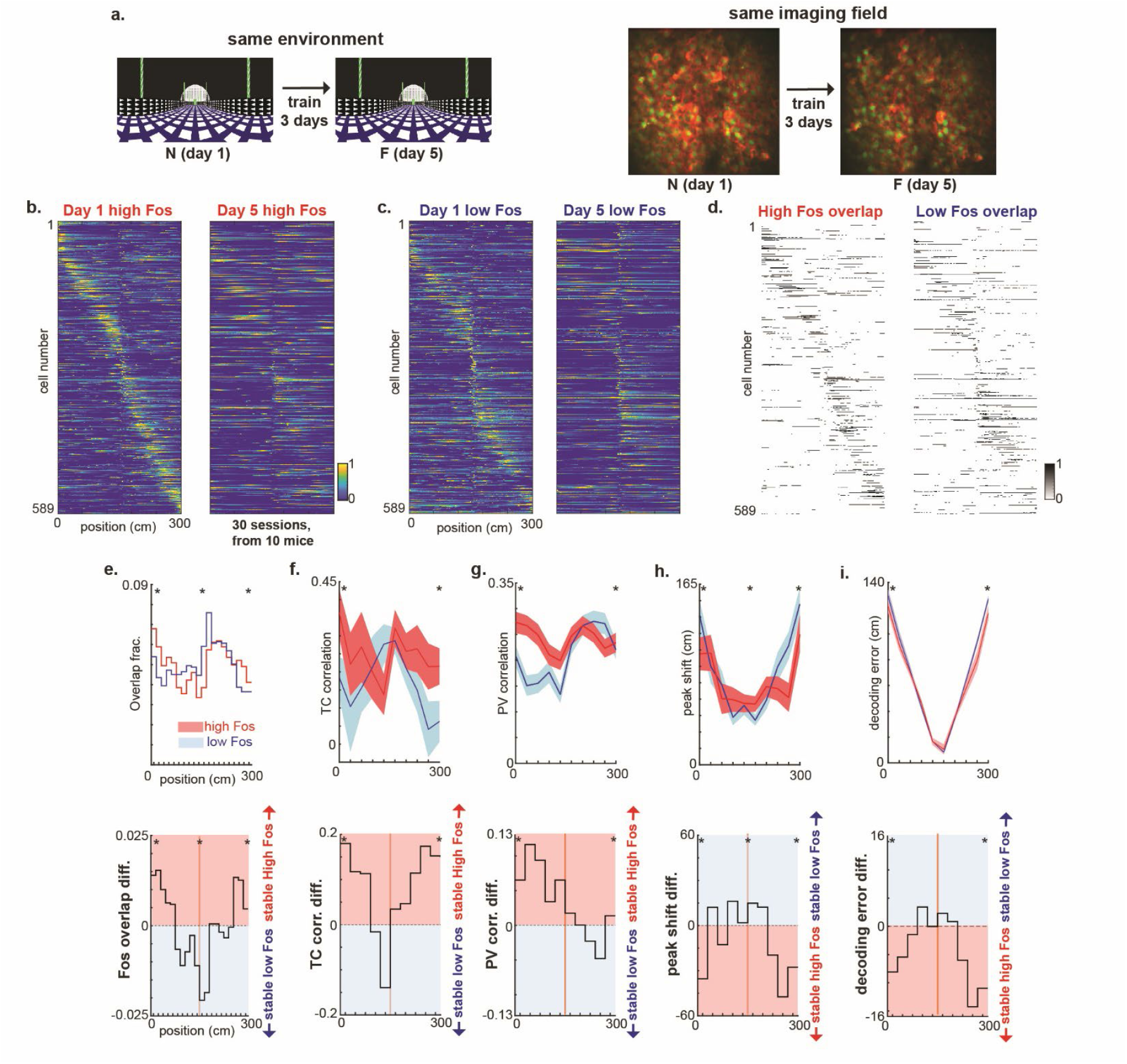
In novel environments, higher *Fos* change predicts stable spatial coding near boundaries across days. **a**. Experimental design. The mice were trained in the same environment for 5 consecutive days (novel on day 1, became familiar over subsequent days). The environment was kept the same, and *Fos* expression change and calcium transients of the cells from the same imaging field were recorded on days 1 and 5 with the same protocol as in Fig. 1. **b-c**. Cross-validated and sorted heatmaps (odd laps sorted by even laps on day 1, all laps plot on day 5, sorted by day 1 even laps) for high *Fos*-change cells (**b**) and low *Fos*-change cells (**c**) on day 1 (left) and day 5 (same cells sorted in same order, right) (n=589 cells for each group, pooled from 30 sessions, 10 mice). **d**. Day 1:day 5 heatmap overlap plots (binary 1 or 0) across days for high *Fos*-change cells (left) and low *Fos*-change cells (right) (n=589 cells for each group, pooled from 30 sessions, 10 mice). **e**. Overlay of high and low *Fos*-change cells overlap (from **d**) fraction across track positions (top) and the difference of high and low *Fos*-change overlap fraction histograms across position bins (bottom, n=10 mice; two-sided Wilcoxon signed-rank test; total 20 position bins, animal-by-animal high vs. low *Fos* boundary zone bins (1-3, 18-20): *P*=1.2e-4, reward zone bins (8-13): *P*=0.025. Star indicates *P*<0.05). **f**. Tuning curve (TC) correlations of high and low *Fos*-change cells across position (top, line indicates average and shaded region indicates s.e.m.) and the difference of high and low *Fos*-change TC correlations across position bins (bottom) (n=331 high *Fos* cells, 353 low *Fos* cells distributed across position; two-sided Wilcoxon rank-sum test; total 10 position bins, high vs. low *Fos* boundary zone bins (1-2, 9-10): *P*=0.032, reward zone bins (4-7): *P*=0.93. Star indicates *P*<0.05). **g**. Population vector (PV) correlations of high and low *Fos*-change cells across position (top, line indicates average and shaded region indicates s.e.m.) and the difference of high and low *Fos*-change PV correlations across position bins (bottom) (n=30 sessions; two-sided Wilcoxon signed-rank test; total 10 position bins, high vs. low *Fos* boundary zone bins (1-2, 9-10): *P*=0.0016, reward zone bins (4-7): *P*=0.40. Star indicates *P*<0.05). **h**. Peak shift of the tuning curve across position (top, line indicates average and shaded region indicates s.e.m.) and the difference of high and low *Fos*-change peak shifts across position bins (bottom) (n=331 high *Fos* cells, 354 low *Fos* cells distributed across position; two-sided Wilcoxon rank-sum test; total 10 position bins, high vs. low *Fos* boundary zone bins (1-2, 9-10): *P*=0.0045, reward zone bins (4-7): *P*=0.0067. Star indicates *P*<0.05). **i**. Bayesian decoding error across position (top, line indicates average and shaded region indicates s.e.m.) and the difference of high and low *Fos*-change decoding error across position bins (bottom) (n=30 sessions; two-sided Wilcoxon signed-rank test; total 10 position bins, high vs. low *Fos* boundary zone bins (1-2, 9-10): *P*=0.0021, reward zone bins (4-7): *P*=0.62. Star indicates *P*<0.05).

*Fos* expression is widely used as a readout of hippocampal engram neurons and has been repeatedly tied to memories of experiences, particularly in contextual fear-conditioning studies^8–10,14,15^. Only recently has a link between *Fos* expression and the hippocampal place code been drawn^11,12^. A previous study^12^ found that low *Fos*-change place cells preferentially encoded reward locations, while stable high *Fos*-change place cells tiled track locations uniformly, thus implying that the engram encodes all spatial locations. However, that study was conducted on a continuous (circular) virtual-reality track that lacked explicit environmental boundaries, yielding a continuous temporal experience for the animal. Here, we introduced a task design that imposed discrete event boundaries using a virtual track with an unambiguous beginning and end. This led to our discovery that high *Fos*-change cells preferentially encode the track boundaries. Thus, when the experience is segmented rather than continuous (e.g., walking down a corridor from one doorway to another, which involves a clear start and an end), high *Fos*-change neurons are more likely to encode the start and end of the episode. This boundary bias adds to the prior observations from continuous environments and suggests that whether high *Fos*-change cells are uniform or boundary-enriched depends on the structure of the experience^16–19^.

Further, we found that novelty-evoked increases in *Fos* expression not only increased the likelihood of neurons becoming place cells, but also predicted the stability of new place cells formed near the boundaries over days^19^. Altogether, these findings present a new model linking place coding and engrams: engram neurons preferentially encode and stabilize the beginnings and ends of new memories, and selective retrieval of these boundaries may initiate recall that propagates to the full experience, possibly via sequence-completion or attractor-like mechanisms^16,20,21^. This boundary emphasis may implement an efficient, segmental code for memory retrieval—assigning greater weight to event boundaries, providing nodes from which the entire memory can be rapidly recalled. This idea also provides a rationale for selective engram reactivation at recall^10,15,22,23^—key segments suffice to recover the whole. Moreover, during representational drift, only a select subset of cells has been observed to remain stable over days^5,6^, particularly those that are highly excitable^6^. Our study suggests that the boundary-stable (more drift-resistant) high *Fos*-change cells could serve as key anchor points from which a complete representation of an experience could be readily reinstated.

Nonetheless, *Fos* expression change is not deterministic—some high *Fos*-change cells encode reward sites and are unstable across days. Thus, *Fos* expression is clearly only part of the multiple molecular pathways connecting gene transcription to spatial coding and memory stability. Future work should examine additional activity-regulated genes (e.g., *Npas4, Arc*, and others^24–26^) to determine the overarching relationship between gene expression change and place cell organization and stability. Additionally, since *Fos* expression ramps up over tens of minutes to hours^12,27^ after the place field formation, it is unlikely that *Fos* itself plays a direct role in the immediate encoding of the boundaries; instead, its primary contribution may be to consolidate and stabilize newly formed memories, potentially via plasticity mechanisms such as bidirectional perisomatic inhibitory plasticity^28^. Future studies can test this causality with targeted *Fos* perturbations (e.g., conditional knockouts)^12,28^.

**Extended Data Fig. 1.**
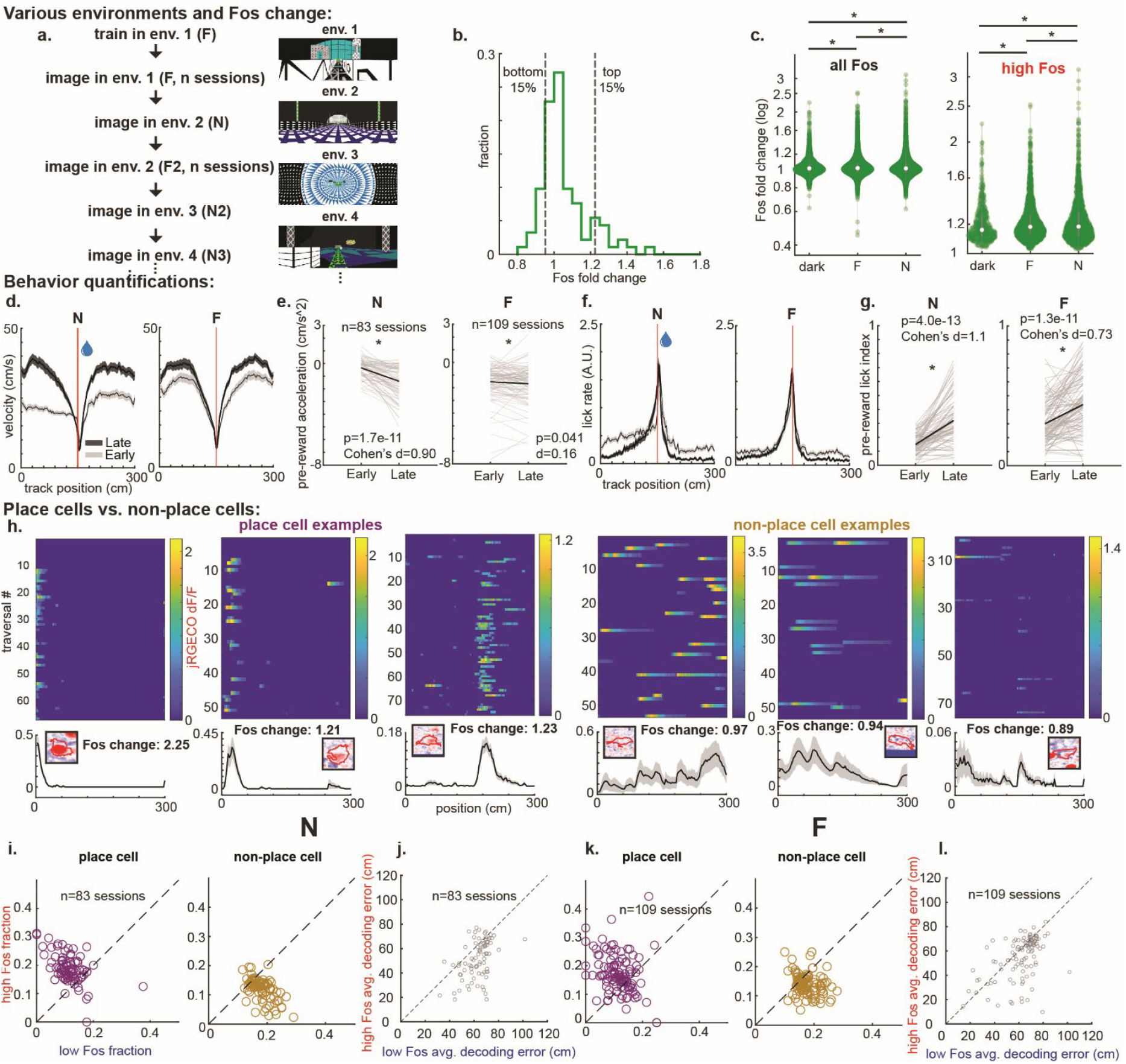
*Fos* fold change in different environments and behavioral quantifications. **a**. Longitudinal experimental design. Mice were exposed to either familiar or novel environments across days. For the environment to be considered familiar (F), mice had to be exposed to the same environment for at least 3 days. **b**. *Fos* fold change histogram of the example imaging field shown in Fig. 1c. The vertical dashed lines indicate the top 15% and bottom 15% of *Fos* fold change. **c**. Cell-by-cell *Fos* fold change for all cells after 3 hrs from the baseline in dark, F, and N environments (left, n=3146 cells in dark, 16117 cells in F, 11637 cells in N; two-sided, two-sample Kolmogorov–Smirnov test; dark vs. F: *P*=0.0042, F vs. N: *P*=3.98e-5, dark vs. N: *P*=0.0046) and for high *Fos*-change cells (right, n=472 cells in dark, 2416 cells in F, 1746 cells in N; two-sided, two-sample Kolmogorov–Smirnov test; dark vs. F: *P*=1.2e-4, F vs. N: *P*=0.0073, dark vs. N: *P*=0.0041). Middle values are medians, and the vertical bars are 25^th^ and 75^th^ percentiles. **d**. Velocity profiles of early and late epochs in N and F. Line indicates average, shaded region indicates s.e.m. (83 sessions from 13 mice for N, 109 sessions from 15 mice for F; two-sided Wilcoxon signed-rank test; N early vs. late mean velocity across the track: *P*=7.3e-14, F early vs. late: 7.3e-8; Cohen’s *D* effect size; N early vs. late: *D*=1.3, F early vs. late: *D*=0.56). **e**. Pre-reward acceleration in N and F. Black lines indicate average, and grey lines indicate individual sessions. (83 sessions from 13 mice for N, 109 sessions from 15 mice for F; two-sided Wilcoxon signed-rank test; N early vs. late pre-reward acceleration: *P*=1.7e-11, F early vs. late: *P*=0.041; Cohen’s *D* effect size; N early vs. late: *D*=0.90, F early vs. late: *D*=0.16). **f**. Lick profiles of early and late epochs in N and F. Line indicates average, shaded region indicates s.e.m. (83 sessions from 13 mice for N, 109 sessions from 15 mice for F; two-sided Wilcoxon signed-rank test; N early vs. late mean lick rate across the track: *P*=5.7e-10, F early vs. late: *P*=1.8e-7; Cohen’s *D* effect size; N early vs. late: *D*=0.84, F early vs. late: *D*=0.48). **g**. Pre-reward lick index in N and F. Black lines indicate average, and grey lines indicate individual sessions. (83 sessions from 13 mice for N, 109 sessions from 15 mice for F; two-sided Wilcoxon signed-rank test; N early vs. late pre-reward lick index: *P*=4.0e-13, F early vs. late: 1.3e-11; Cohen’s D effect size; N early vs. late: *D*=1.1, F early vs. late: *D*=0.73). **h**. Place cell examples (left) and non-place cell examples (right). dF/F vs track position during individual track traversals is shown on top, and the average dF/F is shown on the bottom with associated *Fos* change. The black line indicates the average, and the shaded region the s.e.m (bottom). **i**. The session-by-session N high *Fos*-change fraction of place cells and non-place cells is plotted against the low *Fos*-change fraction. (83 sessions from 13 mice; two-sided Wilcoxon signed-rank test; place cell high *Fos* fraction vs. low *Fos* fraction: *P*=6.4e-10, non-place cell high *Fos* fraction vs. low *Fos* fraction: *P*=1.6e-11; N vs. F place cell - non-place cell high and low *Fos*-change cell fraction difference: *P*=0.036). **j**. The session-by-session N average Bayesian decoding error of high *Fos*-change cells plotted against that of low *Fos*-change cells (83 sessions from 13 mice; two-sided Wilcoxon signed-rank test; high *Fos* vs. low *Fos*: *P*=5.7e-10). **k**. The session-by-session F high *Fos*-change fraction of place cells and non-place cells is plotted against the low *Fos*-change fraction. (109 sessions from 15 mice; two-sided Wilcoxon signed-rank test; place cell high *Fos* fraction vs. low *Fos* fraction: *P*=1.3e-8, non-place cell high *Fos* fraction vs. low *Fos* fraction: *P*=1.3e-9; N vs. F place cell - non-place cell high and low *Fos*-change cell fraction difference: *P*=0.036). **l**. The session-by-session F average Bayesian decoding error of high *Fos*-change cells plotted against that of low *Fos*-change cells (109 sessions from 15 mice; two-sided Wilcoxon signed-rank test; high *Fos* vs. low *Fos*: *P*=5.2e-5).

**Extended Data Fig. 2.**
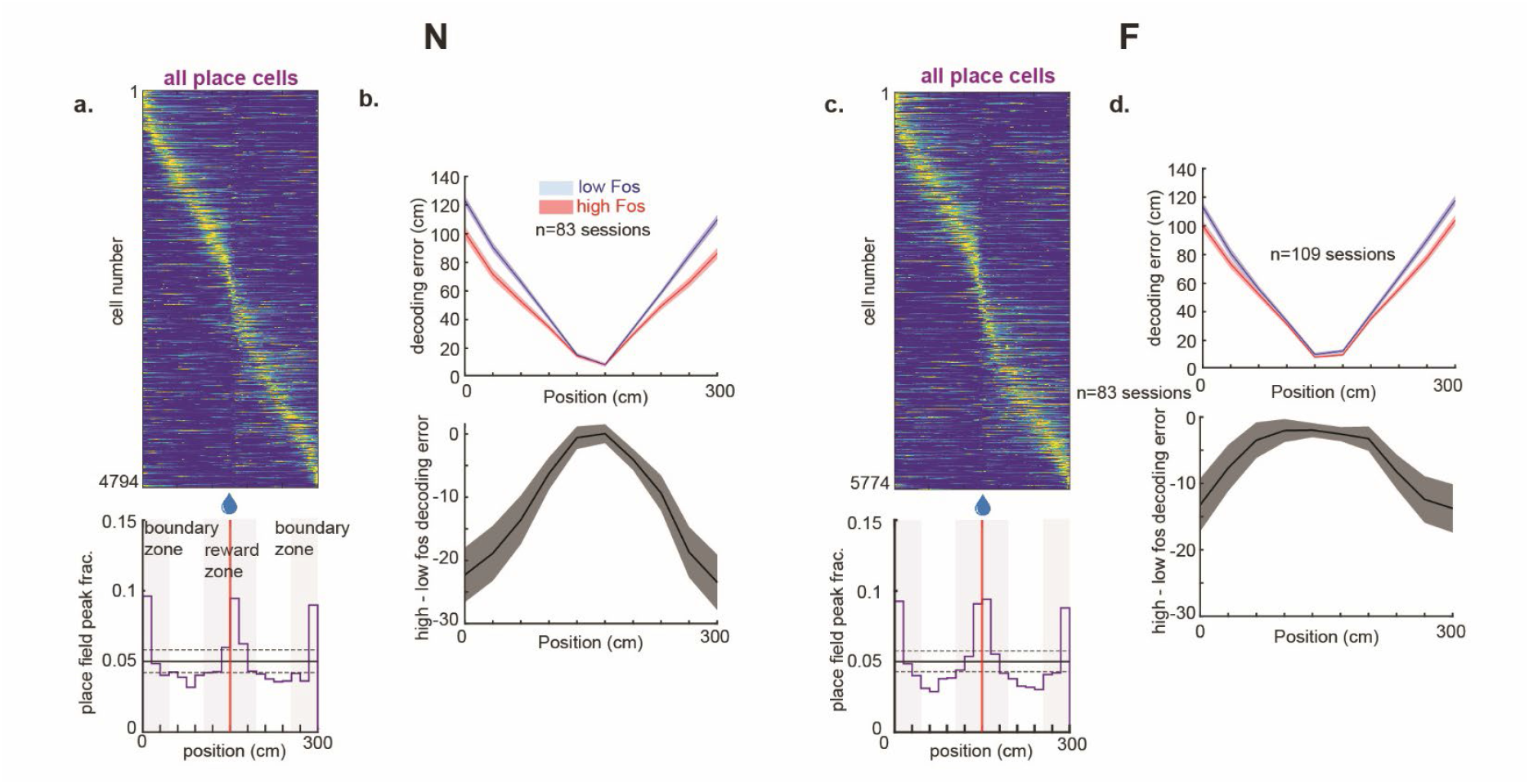
Cross-validated heatmaps of all place cells and Bayesian decoding error of high and low *Fos*-change cells across track position. **a**. Cross-validated and sorted heatmaps (odd laps sorted by even laps, top) for all place cells and their histograms of place field peak fractions (bottom) in N (n=4795 cells pooled from 13 mice). The solid horizontal lines indicate chance level, and dotted horizontal lines indicate two-sided binomial 99% confidence intervals. **b**. The session-by-session N Bayesian decoding error of high and low *Fos*-change cells by track position (top), and difference in decoding error (bottom). The lines are averages and the shaded region indicates s.e.m. (83 sessions from 13 mice; two-sided Wilcoxon signed-rank test; total 10 position bins, high vs. low *Fos* boundary zone bins (1-2, 9-10): *P*=2.6e-21, reward zone bins (4-7): *P*=0.0032). **c**. Cross-validated and sorted heatmaps (odd laps sorted by even laps, top) for all place cells and their histograms of place field peak fractions (bottom) in F (n=5774 cells pooled from 15 mice). The solid horizontal lines indicate chance level, and dotted horizontal lines indicate two-sided binomial 99% confidence intervals. **d**. The session-by-session F Bayesian decoding error of high and low *Fos*-change cells by track position (top), and difference in decoding error (bottom). The lines are averages and the shaded region indicates s.e.m. (109 sessions from 15 mice; two-sided Wilcoxon signed-rank test; total 10 position bins, high vs. low *Fos* boundary zone bins (1-2, 9-10): *P*=1.1e-9, reward zone bins (4-7): *P*=0.0027).

**Extended Data Fig. 3.**
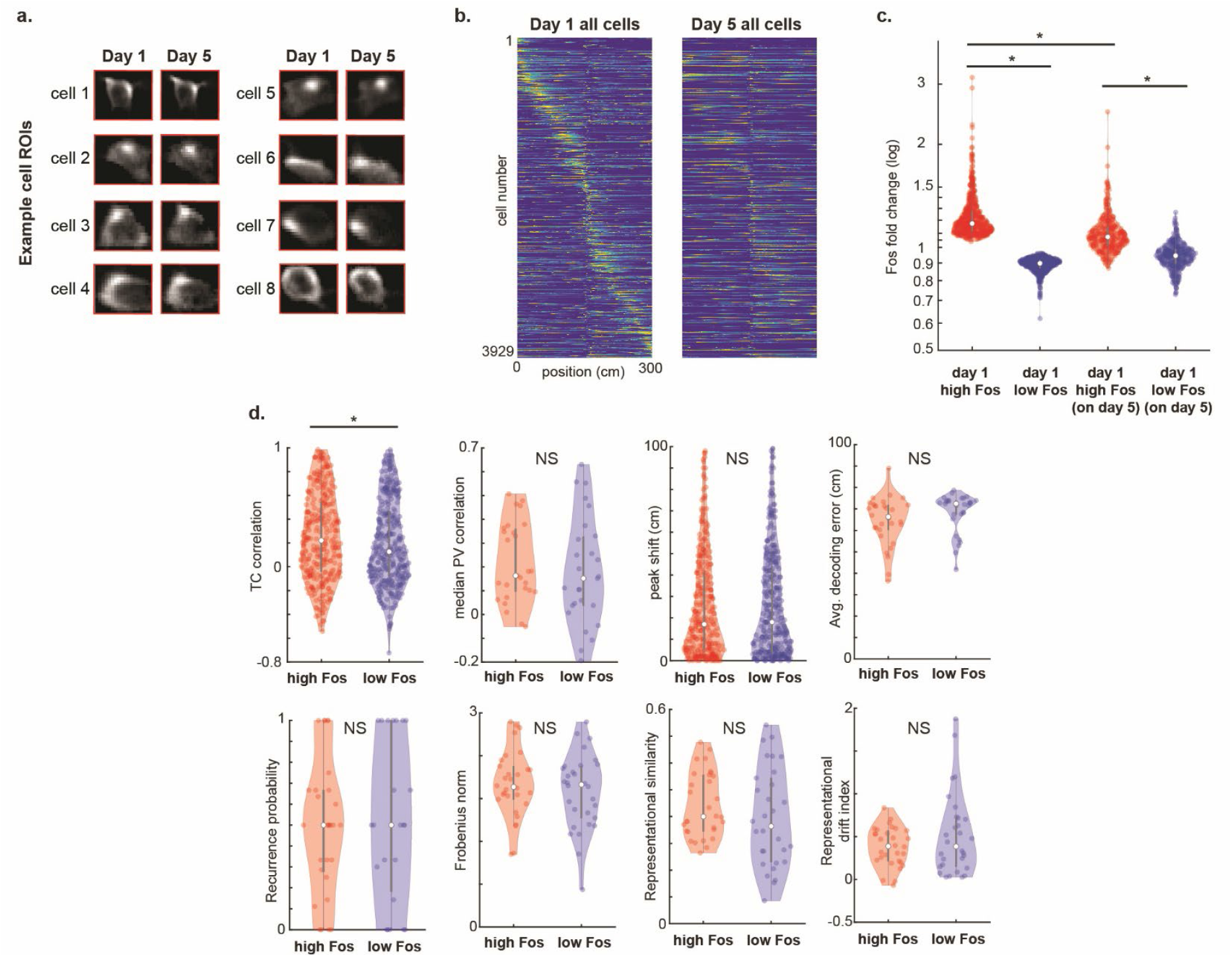
Across-days cell examples and quantifications of representational drift. **a**. Example ROIs identified across days in the imaging field shown in Fig. 3a. Left ROI is from day 1, right ROI is from day 5 (n=8 cells). **b**. Cross-validated and sorted heatmaps (odd laps sorted by even laps on day 1, all laps plotted on day 5, sorted by day 1 even laps) for all cells on day 1 (left) and day 5 (same cells, right). **c**. *Fos* fold change of day 1 high *Fos*-change cells and low *Fos*-change cells on day 5 (n=589 cells for day 1 high and low *Fos* cells, n=331 for day 1 high *Fos* cells on day 5, n=355 for day 1 low *Fos* cells on day 5; two-sided Wilcoxon rank-sum test; day 1 high *Fos* vs. day 1 low *Fos*: *P*=5.5e-194, on day 5 high *Fos* vs. on day 5 low *Fos*: *P*=4.2e-64, day 1 high *Fos* vs. on day 5 high *Fos*: *P*=1.9e-44, day 1 low *Fos* vs. on day 5 low *Fos*: *P*=2.7e-36). **d**. Multiple representational drift measurements of high *Fos*-change cells and low *Fos*-change cells across days. Middle values are medians, and the vertical bars are 25^th^ and 75^th^ percentiles. Star indicates *P*<0.05. NS indicates statistically not significant. TC correlation: n=331 high *Fos* cells, 353 low *Fos* cells, two-sided Wilcoxon rank-sum test, *P*=0.029. PV correlation (median): n=30 sessions, two-sided Wilcoxon signed-rank test, *P*=0.56. Peak shift: n=331 high *Fos* cells, 354 low *Fos* cells, two-sided Wilcoxon rank-sum test, *P*=0.74. Bayesian decoding error (average): n=30 sessions, two-sided Wilcoxon signed-rank test, *P*=0.11. Recurrence probability (average): n=30 sessions, two-sided Wilcoxon signed-rank test, *P*=0.58. Frobenius norm: n=30 sessions, two-sided Wilcoxon signed-rank test, *P*=0.44. Representational similarity: n=30 sessions, two-sided Wilcoxon signed-rank test, *P*=0.18. Representational drift index: n=30 sessions, two-sided Wilcoxon signed-rank test, *P*=0.56.

## Methods

### Mouse strain and surgery

B6.Cg-Tg (Fos-tTA, Fos-shEGFP)1Mmay/J strain from Jackson laboratory (strain# 018306) was used throughout this study to monitor *Fos* expression change (9 male, 6 female adult mice ranging approximately 12 weeks to 16 weeks). For the induction of jRGECO1a in the hippocampus CA1, the mice were anesthetized with 1.5%-2% isoflurane and ~250 nl-310 nl of an adeno-associated virus vector (AAV2/1-CAG-NES-jRGECO1a.WPRE.SV40) with a titer of 1x10e13 genomic copies per ml (GC/ml) was injected using a beveled glass micropipette. A 1 mm diameter craniotomy was performed on the right hippocampus, and the virus was injected at three locations that are ~500-900 µm apart, centered around 1.8 mm lateral, 2.3 mm caudal from bregma, and 1.3 mm beneath the dura surface. After 3-7 days of injection, a hippocampal window was implanted, a stainless-steel cannula with a glass coverslip fixed on one side being placed over the hippocampus. The titanium head plate and light-blocking ring were cemented to the skull afterwards.

### Virtual reality setup and two-photon imaging

Experimental mice were caged singly in identical, unenriched cages (no exercise wheels) in the animal facility and moved to the experimental room in an enclosed, light-proof cart and kept in darkness with broadband white-noise for a minimum of 3 hrs to minimize the effects of external cues before the start of an experiment. Animals were head-fixed on a cylindrical treadmill within a five-monitor virtual-reality (VR) setup with a broadband white-noise during the three hours of the experiment. Visual environments were generated in MATLAB using the ViRMEn engine^29^, and a lick spout was positioned within tongue reach; lick events were detected with a capacitive sensor (SparkFun) and streamed to MATLAB via a National Instruments PCI-6229 DAQ. Visual cues and reward delivery were controlled in MATLAB via the ViRMEn engine. A 4μl water reward was dispensed at 1.5 m (middle) along the 3m track.

Imaging of hippocampal CA1 was performed on a custom movable-objective microscope (Sutter Instruments) with a 40×/0.60 NA air objective (LUCPLFLN40x, Olympus) running ScanImage 5.1. Data was acquired with a unidirectional scanning at 31.25 Hz (512 × 256 pixels). Behavioral signals—including track position, running speed, licking, reward delivery, and per-frame two-photon timestamps—were digitized and synchronized at 1 kHz using a Digidata1440A (Molecular Devices) controlled by Clampex 10.3.

### Training and behavior quantifications

For training, mice were individually housed in identical, unenriched cages (no exercise wheels) and moved to the experimental room in an enclosed, dark cart and kept in the dark with white noise throughout the training sessions; the same was done for the experimental sessions. The mice were water-restricted by receiving a total of 0.8 ml per day and were trained in VR until they displayed anticipatory licking and slowing before the reward.

For behavior quantifications (Extended Data Fig. 1), the session-by-session behavior data were divided into 10 (~3 min each) time bins, and the data within the first bin and last bin were used to compare early vs. late behavior in N and F.

For the session-by-session pre-reward acceleration calculation for each group, the mean acceleration of the mice 30 cm from the reward site was used. For the pre-reward lick index calculation, the sum of licks 30 cm before the reward site was divided by the sum of licks across all track position.

### Acquisition and processing of *Fos* fold change and calcium transients

#### Acquisition for familiar (F) and novel (N) environments

We first obtained baseline time-series in darkness (black screen VR, no rewards) by simultaneously imaging the cfos-shEGFP (920 nm, green) and jRGECO1a (1070 nm, red) channels (1,000 frames within the chosen field). Beginning at VR onset and continuing for 30 min, we recorded 50,000 time-series frames of jRGECO1a calcium transients (920nm laser off). Following the 30 min behavior session, mice remained head fixed in darkness (no VR) without rewards for 2.5 h. We then acquired a z-stack centered on the same imaging field— 25 slices with 2 µm spacing, 200 time-series frames per slice—in both green and red channels for later identification of the same imaging plane in xyz coordinates (in case of any imaging plane drift since baseline time-series acquisition) and compute *Fos* fold change.

#### Acquisition for dark environments

For the dark environment experiments (Extended Data. Fig. 1c), we used the same procedures as the F/N environment experiments, but conducted the entire experiment in darkness and omitted rewards.

#### Extraction of jRGECO1a calcium transients and ROI detection

Registration, ROI segmentation, and raw calcium trace extraction of individual cells were performed using Suite2P^30^ software. Traces were then converted to ΔF/F by an 8th-percentile sliding baseline (500 frames of sliding window). Significant transients were detected based on the ratio of positive versus negative-going transients as described previously^31^.

#### Processing Fos fold change

The image processing pipeline steps used to extract *Fos* fold change are as follows:

1. Extract jRGECO1a (red) and cfos-shEGFP (green) baseline time-series movie from the simultaneously recorded baseline times-series movie (red + green). Motion-correct the jRGECO1a (red) baseline movie (1000 frames, pre-VR) twice using the average jRGECO1a image from Suite2P as a reference image in the xy plane and save the per-frame x,y pixel shifts. Apply these same shifts frame by frame to the cfos-shEGFP (green) baseline movie. Compute the average image of the motion-corrected green baseline movie to obtain the *Fos* baseline reference image.
2. For the post-VR z-stack (25 slices, 200 frames per slice, ~3 hrs after baseline), motion-correct each red z-stack slice twice and save the x,y shifts. Apply the corresponding shifts to the matched green slice. Compute the average image for each motion-corrected green slice.
3. Z-score normalize the green baseline reference image and each z-stack green slice average image (mean 0, s.d. 1). Compute 2D cross-correlation of the normalized baseline reference with each normalized z-series slice to find the slice and x,y pixel offsets that yield the highest correlation peak. Shift that slice by the offsets to overlay it onto the baseline reference in the same x,y,z coordinates.
4. Compute the ratio image as [overlaid green slice (post 3hrs, not normalized) ÷ green baseline reference (before start, not normalized)]. Apply a 2D median filter (51 × 51 px) to the ratio image to estimate the background. Divide the original ratio image by this filtered image to perform background correction. The resulting image is the cfos-shEGFP fold change image (Fig. 1c).
5. Using the jRGECO1a cell ROIs from Suite2P, identify the mean pixel values of the cfos-shEGFP fold change image. This is the *Fos* fold change value for each cell ROI (Fig. 1d, Extended Data Fig. 1b-c).

### Identification of the same cells across days

#### Volumetric plane registration

For high precision, quantitative identification of the same cells on day 1 and day 5, we utilized the online (pre-experiment) volumetric plane registration method as described previously^6^. In brief, on day 1, we attained 1000 frames of the imaging field in the red channel (jRGECO1a), motion corrected, and averaged to attain the target image. Before the start of day 5 experiment, we acquired a 15-slice z-stack (100-200 frames/slice) at 2 µm spacing around the putative target plane. Each slice was motion-corrected and averaged, z-score normalized, and 2D cross-correlated with the z-score normalized day 1 target image. The slice and x,y offsets that maximized the peak correlation defined the best-matching xyz position (offsets reported in µm). We used these offsets to fine-position the objective and precisely and reliably return to the same plane after five days (Fig. 3a, Extended Data Fig. 3a).

#### ROI detection across days

CellReg^32^ was used to align ROIs that were identified across days. To ensure the accuracy of the matching ROIs across days, every individual ROIs’ cellular and subcellular morphology was visually inspected (Fig. 1c-d, Extended Data Fig. 3a).

### High *Fos*-change cells and low *Fos*-change cells

Across all sessions, we excluded cells whose *Fos* change could not be computed because of lateral (x–y) drift of the imaging field after 3 hrs. We also excluded ROIs in a peripheral margin to avoid potential edge-related low signal-to-noise ratio (SNR): cells within 1/14 of the field dimension from any edge were removed. For the top 10% of sessions showing the largest center–edge disparity in *Fos* fold change (quantified by ROI-centroid distance from the field center), we applied a stricter exclusion, removing cells within 1/7 of the field edges.

We then categorized cells into high and low *Fos* change cells using distributional cutoffs from MATLAB’s *quantile* function. We compiled the *Fos* change metric per cell and computed the 15th and 85th percentiles (Extended Data Fig. 1b). Cells with values ≥ 85th percentile were labeled high *Fos*-change, those ≤ 15th percentile were labeled low *Fos*-change, and all others were unclassified. When many cells were tied exactly at a threshold, we included/excluded ties to keep groups exactly at 15% per session; thresholds were independently computed within each session.

### Place cells and non-place cells

Using time epochs during movement (speed ≥ 1 cm s^−1^), spatial information (bits/action potential) was calculated from the following formula as described previously^33^:

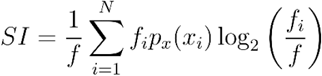

*SI* is the spatial information (bits per action potential), *N* is the total number of spatial bins, *f*_*i*_ is the mean fluorescence in bin *i* is the overall mean fluorescence across all bins, and *p*_*x*_ (*x*_*i*_) is the occupancy probability of the bin *i*.

We built a null distribution by randomly shifting the fluorescence trace with respect to track position and recalculating tuning maps 1,000 times. Cells were considered place cells when their information surpassed 99% of shuffles and had information ≥0.5 bits per action potential. A cell that was not categorized as a place cell was classified as a non-place cell (Extended Data Fig. 1h, binary classification).

### Bayesian decoding error of position

We decoded position in 0.1s time bins (*δt*) using a naïve Bayes model on time epochs only during movement (speed ≥ 1 cm s^−1^). The track was divided into 60 spatial bins, and for each 0.1s time bin, we estimated the conditional probability that the animal occupied a spatial bin *x*_*i*_, given the observed significant transients using the formula:

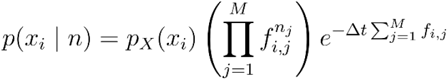

*p*(*x*_*i*_ |*n*) is the probability of occupancy in the bin *x*_*i*_ during a time window n, *f*_*i,j*_ is the average rate of significant transients of a neuron *j* in bin *x*_*i*_, and *M* is the total number of neurons. The spatial bin with the highest posterior probability was taken as the decoded position.

For within-day population decoding of the position of high *Fos*-change cells and low *Fos*-change cells (Extended Data Fig. 1j,l,2b,d), we used the same number of high *Fos*-change cells as low *Fos*-change cells within each session (because groups were defined by quantiles within each session, equal numbers of high and low *Fos*-change cells were ensured a priori). For each session, the entire session was divided into individual laps, and the position decoding for each lap (test data set) was done by allocating all other laps as the train data set. This was iterated over all the laps to test the decoding error (difference between decoded position and real position) for each lap, and the average decoding error per spatial bin was attained per session afterwards.

### Reward zone vs. boundary zone quantifications

Track position was divided into 100 bins, and each neuron’s average transient peak bin location was extracted. For the reward and boundary zone fraction calculations (Fig. 2), we calculated the fraction of place field peaks in each zone after pooling cells across each animal’s sessions, repeating the calculation as the zone window was broadened from 1 to 15 bins. We took the average across all zone sizes (1 to 15 bins) to attain the average reward zone and boundary zone fraction values for comparing high *Fos*-change cells and low *Fos*-change cells.

For quantifications of reward zone and boundary zone fractions across time (Fig. 2 k,l), sessions were segmented into 10 (~3 min) time bins. Within each animal, high/low *Fos*-change cells were pooled, transients were mapped to 100 spatial bins, and the fraction of peak bins within the reward and boundary zones was computed as the zone width was broadened from 1 to 15 bins. We then averaged across zone widths (1 to 15 bins) to obtain the reported fractions for each time bin. For quantifications of the change of reward or boundary zone fractions over time bins, we fitted regression lines as a function of time bins (animal-by-animal) and quantified whether the distributions of the slopes of the regression lines were different from zero (one-sample, two-sided t-test).

For reward and boundary zone statistics in Fig. 3e overlap fractions, we pooled over 3 bins at the start/end (total 6 bins) and 6 bins at the reward (out of a total of 20 bins). For Fig. 3f-i metrics, we pooled over 2 bins at the start/end (total 4 bins) and 4 bins at the reward (out of a total of 10 bins).

### Spatial overlap and representational drift measurements across days

#### Spatial overlap

We quantified day 1 to day 5 spatial overlap by first binarizing (1 or 0) spatially binned (100 bins), normalized activity maps on day 1 (cross-validated odd laps sorted by even laps) and day 5 (all laps, Fig. 3b,c) with a threshold of 0.3, yielding 0 or 1 for each cell *C* at spatial bin *i*. The overlap (Fig. 3d) at the spatial bin *i* for each cell was 1 only if both day 1 and day 5 spatially binned transients at that bin were 1.

We then calculated the overlap fraction at the spatial bin *k* (out of 20 bins) by summing all values of 1 across all sorted cells in the bin *k* and dividing by the sum of all 1s across all *k* bins.

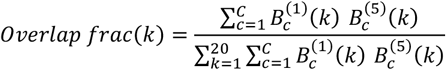

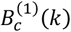 and 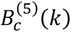 are day 1 and day 5 binary values (1 or 0) at bin *k* for cell *C*.

#### Tuning curve (TC) correlation

After cells were binned into 100 spatial bins, the tuning curve vector correlations (Pearson correlation of the same row x row of heatmaps) of cells that were active on both days were calculated. High and low *Fos* change cells that were inactive on either day were excluded. For TC correlation calculation across positions, cells were assigned to a position bin based on their peak tuning curve location on day 1.

#### Population vector (PV) correlation

After cells on day 1 and 5 were binned into 100 spatial bins and sorted by their peak bin location on day 1, the population vector correlations (Pearson correlation of the same column x column of heatmaps) were computed bin by bin. All cells in high and low *Fos* change groups were used, and for the cells that were inactive on one of the days, the data were filled with 0s.

#### Peak shift

After cells were binned into 100 spatial bins, their day 1 and day 5 tuning curve peak bin locations were identified, and the absolute value of their differences was calculated. High and low *Fos* change cells that were inactive on either day were excluded.

#### Bayesian decoding error

For decoding across days, we used all high and low *Fos* change cells that were active on day 1 (same number of cells in each group), using the calculation as mentioned above. For cells that were inactive on day 5, data were filled with 0s.

#### Recurrence probability

A cell was labeled recurrent (1) if it (i) was active on both days in >1/3 of laps and (ii) had a cross-day tuning-curve (TC) correlation > 0.38; otherwise it was labeled 0. The 0.38 cutoff was derived from a one-predictor logistic regression trained to distinguish same-cell odd–even laps within-day TC correlations from a 1,000× cell-identity–shuffled within-day null (day 1, all laps), using the p=0.5 decision boundary.

#### Frobenius norm

The peak-normalized spatial maps (day 1 and day 5) were used to calculate the session-by-session Frobenius norm, each session normalized by the square root of the number of cells.

#### Representational similarity

The peak-normalized spatial maps (day 1 and day 5) were linearized, and correlation coefficients were acquired to assess representational similarity^34^. For cells that were inactive on day 5, data were filled with 0s before the correlation.

#### Representational drift index

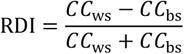

*CC*_*ws*_ was defined as the linearized correlation coefficient of within-session even and odd laps of peak-normalized spatial maps on day 1, and *CC*_*bs*_ was defined as the linearized correlation coefficient across days 1 and 5 (all laps) to assess the representational drift index^34^.

